# Optimization of Tenocyte Lineage-related Factors from Tonsil-derived Mesenchymal Stem Cells using Response Surface Methodology

**DOI:** 10.1101/327734

**Authors:** Seung Yeol Lee, Hyang Kim, Sang-Jin Shin, Soon-Sun Kwon

## Abstract

Researchers should consider various potential factors that affect tenogenic differentiation of mesenchymal stem cells (MSCs); however, this requires numerous experimental settings, which are associated with high cost and time. We aimed to assess the differential effects of transforming growth factor beta 3 (TGF-β3) on the tenogenesis of tonsil-derived MSCs (T-MSCs) and bone marrow-derived MSCs (BM-MSCs) using design of experiments (DoE). Bone marrow and tonsillar tissue was collected from four patients; mononuclear cells were separated and treated with 5 and 10 ng/mL of TGF-β3 with vehicle control. A full-factorial experimental design with a categorical factor of 0 was employed to study the effect of tension based on T-MSCs. Eighty-four trials were utilized, fitted with RSM, and then used to obtain mathematical prediction models. Exposure of T-MSCs and BM-MSCs to TGF-β3 increased the expression of scleraxis (SCX), tenomodulin (TNMD), decorin, collagen I, and tenascin C. Expression of most of these factors reached a maxima after 2–3 days of treatment. Considering all of the tenocyte lineage-related factors that were assessed, the predicted value of the factors from T-MSCs was significantly induced at 2.7 ng/mL of TGF-β3 during 2.5-day culture, whereas the predicted value of the factors from BM-MSCs was significantly induced during 2.3-day culture, regardless of TGF-β3 concentration. This study demonstrated that tenogenic differentiation of T-MSCs and BM-MSCs under TGF-β3 stimulation showed a similar culture time for peak expression of tenocyte-related mRNAs using RSM. This study suggests the potential of using the DoE approach for optimization of the culture protocol for tenogenesis of MSCs.

## Introduction

Tissue healing after tendon repair surgery is challenging [1] because tendons have limited self-healing potential given the scar tissue on the repaired tendon [2, 3]. Even after long periods, the structure and strength of the repaired tissue do not show full recovery, and the tissue does not return to the normal state [4]. For these reasons, various studies on tendon repair using tissue engineering with mesenchymal stem cells (MSCs) have been performed [5–9].

Although bone marrow-derived, adipose tissue-derived, synovial membrane-derived, and human embryonic MSCs have been used to differentiate into tenocytes [6, 9, 10], these stem cells are obtained through invasive procedures. Thus, there is usually a lack of adequate number of MSCs for clinical use [11]. T-MSCs obtained from waste tissue after tonsillectomy represent a new source of progenitor cells [12, 13], and several studies have focused on T-MSCs as cellular therapeutic agents for various diseases. If T-MSCs show non-inferior tenogenic differentiation potential compared to other cell sources, physicians can consider tonsils as a stem cell source for cellular therapeutic agents. However, protocols for tenogenic differentiation of T-MSCs have not been established, and there is a lack of studies comparing T-MSCs with MSCs from other cell sources for tenogenic differentiation.

Researchers should consider various potential factors affecting tenogenic differentiation of MSCs; however, this requires numerous experimental settings whose optimization is associated with high cost and time. Response surface methodology (RSM), which is a part of design of experiments (DoE), is gaining recognition as a powerful approach for optimizing conditions for the production of industrially important products such as chemicals and enzymes. In the last few years, RSM has been applied to optimize and evaluate the interactive effects of independent factors in numerous chemical and biochemical processes [14]. Recently, DoE has been used to investigate differentiation of MSCs [15]. The main advantages of this methodology are that it (1) avoids experimental bias and (2) reduces the number of experiments, leading to a rational understanding of what could be the most favorable factor combination [14, 16].

In this study, we aimed to optimize culture conditions for the tenogenesis of T-MSCs and BM-MSCs using TGF-β3 at different concentrations and culture times.

## Materials and methods

This experimental study was approved by the institutional review board of our institution. Informed consent was obtained from all of the patients or patients’ legal guardians.

### Isolation and cultivation of human MSCs

Bone marrow was collected from four patients (mean age: 79.0±2.2 years) and mononuclear cells were separated using the Ficoll-Paque Premium (GE Healthcare, Chicago, IL, USA) gradient method. The isolated cells were seeded at a density of 1×10^5^ cells/cm^2^ in a growth medium consisting of low-glucose Dulbecco’s Modified Eagle’s Medium (DMEM-LG; Hyclone, South Logan, UT, USA), 10% fetal bovine serum (FBS; Corning, Corning, NY, USA), 100 U/mL penicillin, and 100 μg/mL streptomycin. The tonsillar tissues were collected from four patients (mean age: 7.6±0.6 years) and minced by using surgical scissors, followed by enzymatic digestion using 210 U/mL collagenase type I (Sigma, St. Louis, MO, USA) and 90 KU/mL DNase (Sigma, St. Louis, MO, USA) in DMEM-LG for 30 min at 37°C. Following filtration through a 100-μm cell strainer (BD Bioscience, Franklin Lakes, NJ, USA), the cells were washed twice with Dulbecco’s phosphate-buffered saline (D-PBS; Chembio, Seoul, Korea) Mononuclear cells were separated using the Ficoll-Paque gradient method [13]. The cells were seeded at a density of 1×10^4^ cells/cm^2^ in growth medium. MSCs were incubated in a 5% CO_2_ incubator with humidified air at 37°C, and the medium was replaced every other day. After reaching 80% confluency, the cells were split at a ratio of 1:3 for BM-MSCs and 1:4 for T-MSCs. MSCs were used between passages 2 and 4 for further experiments. Flow cytometry was performed to characterize the immunophenotype of BM-MSCs and T-MSCs using the human MSCs analysis kit (BD Bioscience, Franklin Lakes, NJ, USA).

### Tenogenic differentiation of BM-MSCs and T-MSCs

BM-MSCs and T-MSCs were seeded at a density of 1 × 10^4^ cells/cm^2^ into 24-well plates in growth medium. After 18 h, the growth medium was removed and replaced with tenogenic differentiation media, which consisted of DMEM-LG, 10% FBS, and 50 μg/mL L-ascorbate-2-phosphate with 5 or 10 ng/mL TGF-β3 (Sigma, St. Louis, MO, USA). MSC growth medium was added to the control group. The medium was replaced three times a week.

### Isolation of RNA and quantitative real-time PCR

Total RNA was isolated daily for up to 7 days using the total RNA mini kit supplemented with DNase I (NucleoGen Biotechnology). First-strand cDNA was synthesized using SuperScript III First-Strand cDNA synthesis kit (Invitrogen), and quantitative real-time PCR was performed with SensiFAST SYBR Hi-ROX kit (Bioline) using the QuantStudio 3 real-time PCR system (Applied Biosystem, Life Technologies). The relative expression level of each gene was normalized to that of 18S rRNA and calculated using the 2^−ΔΔCt^ method. The data are presented as fold changes relative to controls. The following genes were analyzed: scleraxis (SCX), tenomodulin (TNMD), decorin, collagen I/III, and tenascin C. These genes have a crucial role in the tenogenesis of MSCs [17–19]. The primers used in this study are shown in Table 1.

**Table 1.**
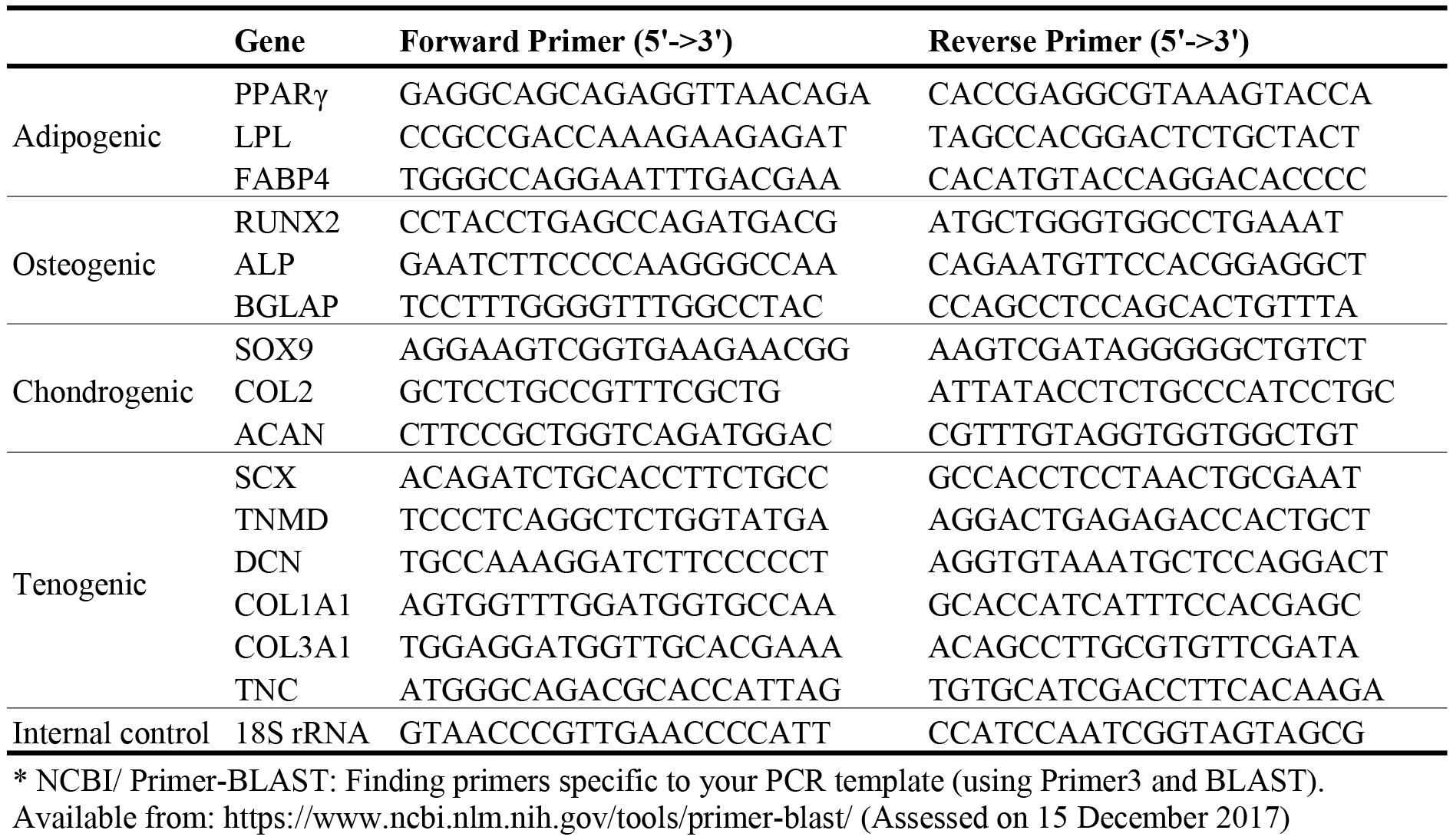
Primers used in this study.*

### Experimental design

A full-factorial experimental design with a categorical factor of 0 was employed to study the effect of tension in T-MSCs. The design comprised of three levels coded as −1, 0, and +1. In total, 18 runs were performed in duplicate to optimize the level of chosen variables, such as days and amount. For the purpose of statistical computation, two independent variables denoted as *x*_1_ and *x*_2_ were selected. The levels used in the experiments were determined from the preliminary experiments and are presented in Table 2.

**Table 2.** ANOVA for response surface model

| MSCs | Source | Sum of squares | Mean square | F-value | p-value |
| --- | --- | --- | --- | --- | --- |
| T-MSCs | Days | 17.57867 | 5.859557 | 9.98 | < 0.001 |
| | Concentration of TGF-β3 | 88.30324 | 29.43411 | 50.11 | < 0.001 |
| BM-MSCs | Days | 37.2556 | 12.4185 | 3.68 | 0.013 |
| | Concentration of TGF-β3 | 71.4878 | 23.8293 | 7.07 | < 0.001 |

MSCs, mesenchymal stem cells; T-MSCs, tonsil-derived mesenchymal stem cells; BM-MSCs, bone marrow-derived mesenchymal stem cells

The results were analyzed via analysis of variance (ANOVA). Multi-level factorial designs were used to estimate the response calculated according to the following second-degree polynomial equation (1):

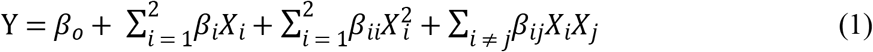

where Y is the estimated response; *β*_*o*_, *β*_*i*_, *β*_*ii*_, and *β*_*ij*_ are the regression coefficients for intercept, linearity, square, and interaction, respectively; and *X*_*i*_ and *X*_j_(i,j = 1,2, i ≠ j) are the different interaction coefficients between the predicted response and independent variables in the coded values according to Table 2.

In this study, RSM combined with a full factorial design was used to investigate MSCs in tissues. By using a multi-level two-factorial design and a full range of RSM, the following parameters were optimized: SCX, TNMD, decorin, collagen I/III ratio, and tenascin C. A second-order polynomial regression model was used to generate three-dimensional response surfaces of MSCs. The regression model would provide a good explanation of the relationship between the independent variables and responses [20].

### Statistical analysis

The statistical significance among different concentrations and time points was analyzed by ANOVA, followed by Tukey’s multiple comparison test using SAS version 9.4.2 (SAS Institute, Cary, NC). All statistics were two-tailed, and p < 0.05 was considered significant.

## Results

### Immunophenotypic characterization of BM-MSCs and T-MSCs

In this study, immunophenotypic surface marker analysis of BM-MSCs and T-MSCS showed the typical expression profile of human MSCs. Both MSC populations expressed CD73, CD90, and CD105, whereas they lacked expression of CD11b, CD19, CD34, CD45, and HLA-DR (Table 3). T-MSCs showed stemness and were able to differentiate into at least adipocytes, osteoblasts, and chondroblasts.[21]

**Table 3.**
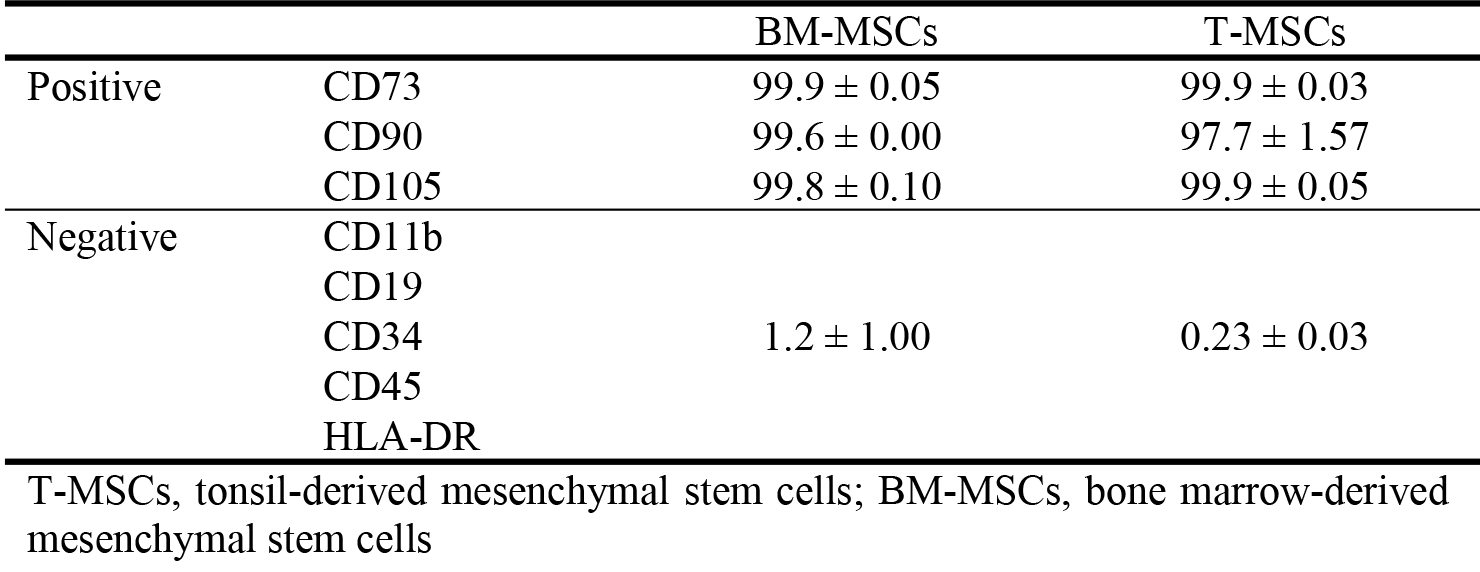
Immunophenotypic markers on the cell surface of T-MSCs and T-MSCs.

### Expression of tenogenic genes in MSCs under TGF-β3 stimulation

Exposure of T-MSCs and BM-MSCs to TGF-β3 resulted in increased expression of SCX, TNMD, and tenascin C, as well as increased collagen I/III ratio. The expression of decorin in both T-MSCs and BM-MSCs was lower than that of untreated MSCs. The peak expression of each mRNA varied slightly according to the culture time and TGF-β3 concentration (Fig 1).

**Fig 1.**
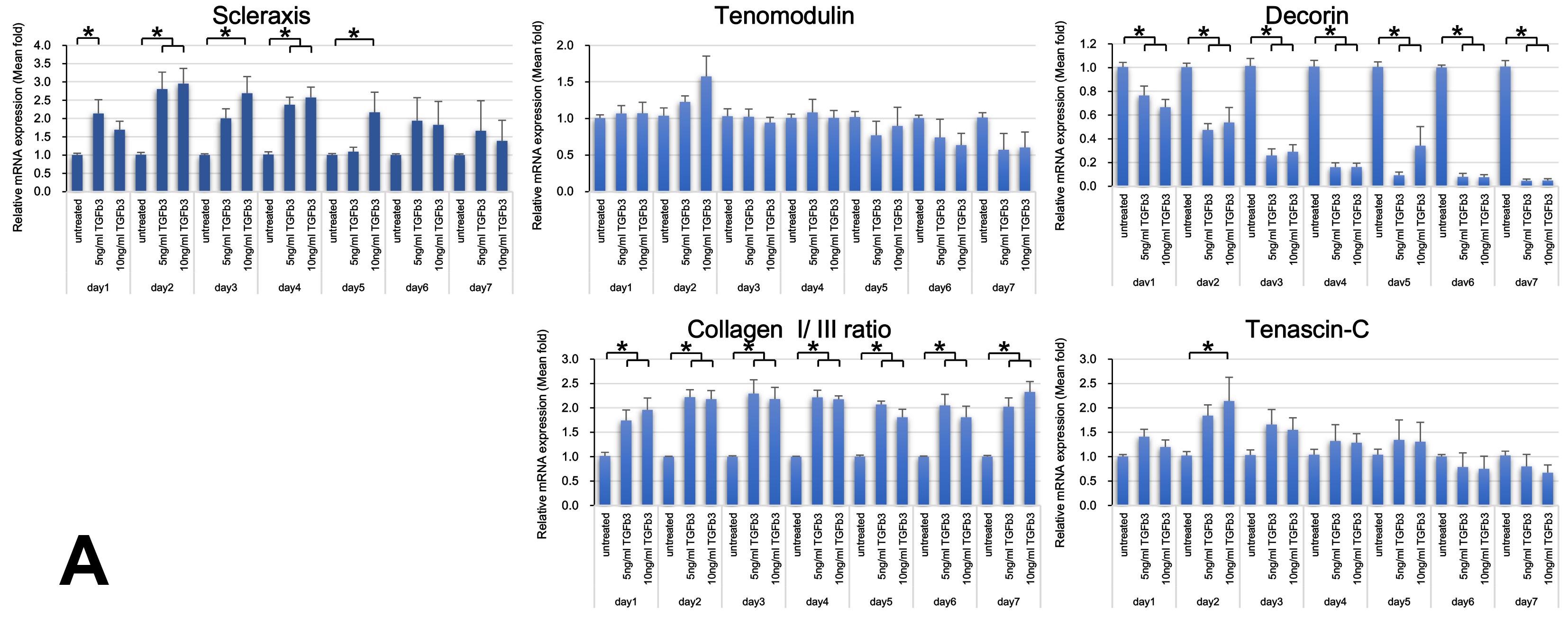
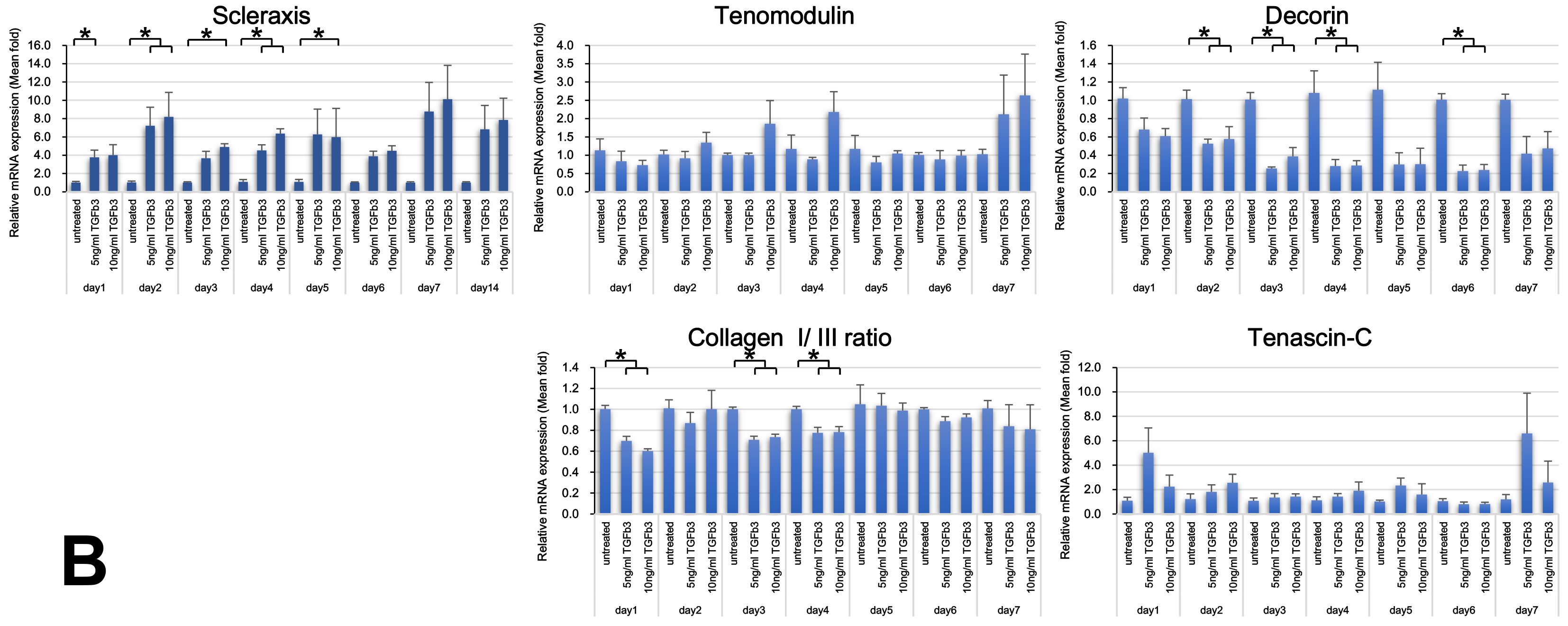
Tenogenic mRNA expression of T-MSCs (A) and BM-MSCs (B) exposed to TGF-β3. The peak expression of each mRNA differed slightly according to the culture time and TGF-β3 concentration. The collagen I to III ratio of T-MSCs increased regardless of TGF-β3 concentration, whereas the ratio in BM-MSCs decreased. (*; p < 0.05, analyzed by one-way analysis of variance, followed by Tukey’s multiple comparison test)

### Optimization of tenocyte lineage-related factors from T-MSCs

The DoE used in this study allowed optimization of tenocyte lineage-related factors from T-MSCs and BM-MSCs under different TGF-β3 concentrations and culture times. From the DoE approach, the predicted value of SCX from T-MSCs was significantly increased at 8.4 ng/mL TGF-β3 (p = 0.014) and 2.3 days of culture (p = 0.040). The expression of collagen I showed the maximum increase at 8.1 ng/mL TGF-β3 (p < 0.001) at 2.7 days of culture (p = 0.036). TNMD peaked at 2.5 days of culture (p = 0.011), regardless of TGF-β3 concentration. TGF-β3 concentration affected the peak expression of decorin (p < 0.001) and the ratio of collagen I to III (p < 0.001) regardless of culture time (Table 4). For all of the tenocyte lineage-related mRNAs that were assessed, the predicted value of the factors was significantly increased at 2.7 ng/mL TGF-β3 (p < 0.001) at 2.5 days of culture (p = 0.001) (Fig 2A).

**Table 4.**
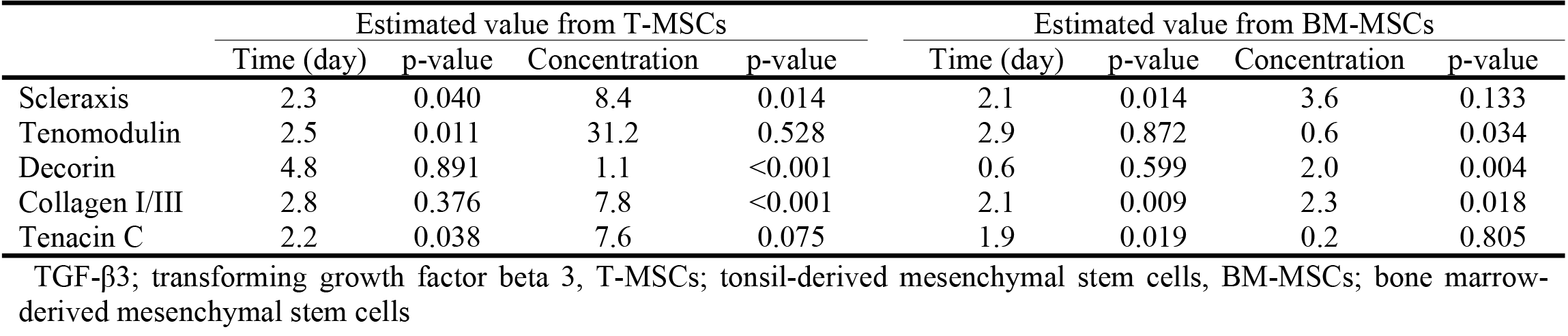
Optimization of each tenocyte lineage-related factor from T-MSCs and BM-MSCs using response surface methodology.

**Fig 2.**
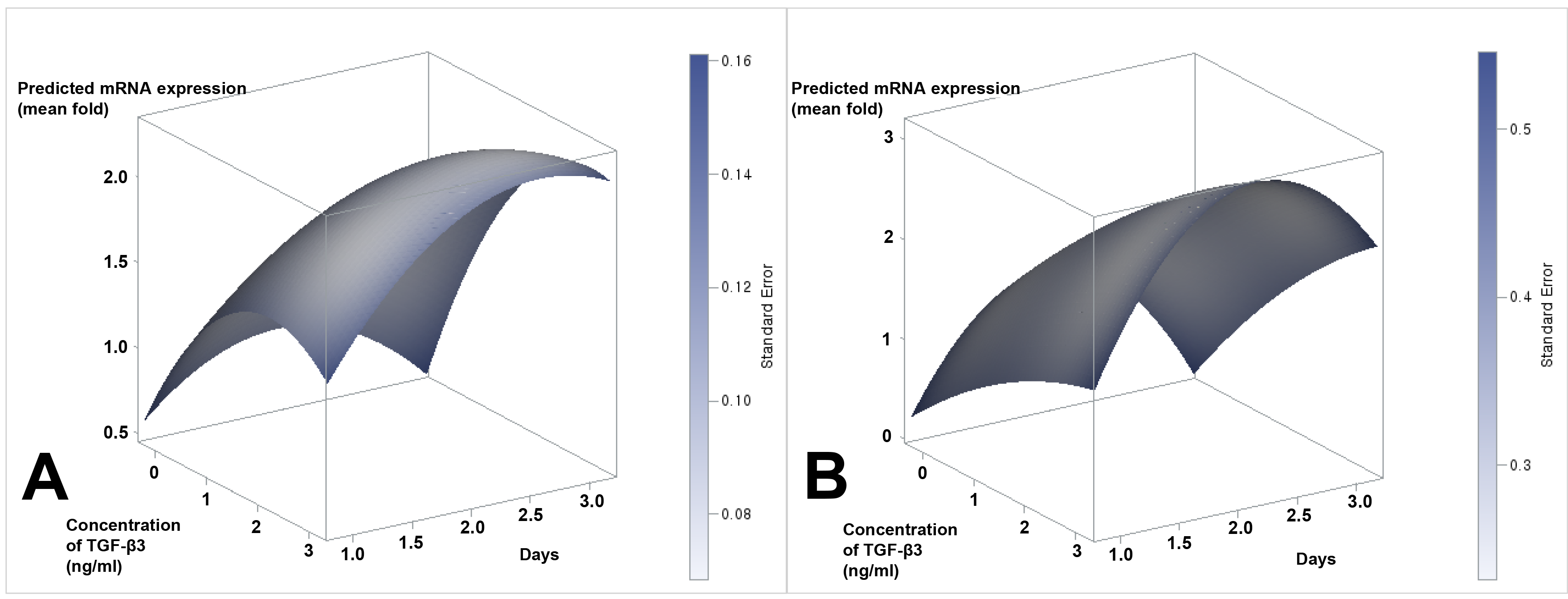
Response surface for all tenocyte lineage-related mRNAs from T-MSCs (A) and BM-MSCs (B).

### Optimization of tenocyte lineage-related factors from BM-MSCs

From the DoE approach, the predicted maximum ratio of collagen I to III in BM-MSCs was significantly increased at 2.3 ng/mL TGF-β3 (p = 0.018) and 2.1 days of culture (p = 0.009). The maximum expression of SCX, collagen I, and tenascin C was affected by culture time. The expression of TNMD and decorin peaked at 0.6 ng/mL (p = 0.036) and 2.0 ng/mL (p = 0.004) TGF-β3, respectively, regardless of culture time for the predicted peak expression of TNMD (p = 0.872) and decorin (p = 0.599) (Table 4). For all of the tenocyte lineage-related mRNAs that were assessed, the predicted value of the factors was significantly increased at 2.3 days of culture (p = 0. 004) regardless of TGF-β3 concentration (Fig 2B).

## Discussion

In this study, we aimed to assess the effects of TGF-β3 on the tenogenesis of T-MSCs and BM-MSCs using RSM and found that tenocyte-like cells could be successfully differentiated from T-MSCs and BM-MSCs under TGF-β3 stimulation.

During tenogenic differentiation of T-MSCs, researchers should consider various potential factors influencing tenogenesis; however, this would require numerous experimental settings. We conducted this study to optimize the culture conditions for tenogenesis of T-MSCs and BM-MSCs under TGF-β3 stimulation at different concentrations and culture times using design experiments. In the present study, exposure of T-MSCs and BM-MSCs to various concentrations of TGF-β3 resulted in an increase in the expression of SCX, TNMD, decorin, collagen I, and tenascin C after 2–3 days of culture. Considering all of the tenocyte lineage-related factors that were assessed, the predicted value of the factors was significantly induced during 2.5-day culture and 2.3-day culture in T-MSCs and BM-MSCs, respectively.

In the present study, the expression of scleraxis, TNMD, collagen I/III ratio, and tenascin C, i.e., all examined molecules except for decorin, tended to increase after exposure of T-MSCs and BM-MSCs to TGF-β3. Our results were similar to those of a previous study [19]. Although the peak expression of each gene for T-MSCs and BM-MSCs was analyzed using DoE, the optimal culture time and concentration of TGF-β3 varied for the peak expression of each gene. For instance, in the present study, the expression of decorin decreased as the culture progressed, whereas the expression of TNMD peaked during the 7-day culture in BM-MSCs after TGF-β3 treatment. Because the expression levels of all genes that were analyzed in this study should increase for tenogenesis of MSCs, we could not determine the optimal culture time. Figure 1 shows the results of ANOVA for mRNA expression. Although the expression of TNMD did not change significantly, we could not determine the expression of TNMD after exposure of MSCs to TGF-β3 owing to the small sample size and experimental settings. A full factorial design is a powerful tool for understanding complex processes of tenocyte lineage-related factors in multifactor systems, because it includes all possible factor combinations for each of the factors. RSM is an empirical statistical technique employed for multiple regression analysis that uses quantitative data obtained from design experiments to solve multivariate equations simultaneously.

### Statistical analysis

The quadratic equation for predicting the optimum point was obtained according to the data and input variables, and then the empirical relationship between the response and independent variables in the coded units was presented based on the experimental results as follows:

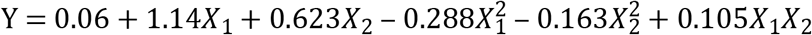

where Y is the T-MSCs, and *X_1_ and X_2_* are the day and amount, respectively (Table 5). The results of the ANOVA for the quadratic equation are tabulated in Table 5. The ANOVA shows whether the equation and actual relationship between response and significant variables represented by the equation are accurate. The significance of the coefficient term is determined by the values of *F* and *p*, and larger values of *F* and smaller values of *p* represent more significant terms.

**Table 5.** ANOVA for response surface quadratic model

| MSCs | Source | Estimate | Standard Error | F-value | p-value |
| --- | --- | --- | --- | --- | --- |
| T-MSCs | Intercept | 0.064 | 0.337 | 0.04 | 0.849 |
| | $X_1$ | 1.143 | 0.352 | 10.50 | 0.0013 |
| | $X_2$ | 0.623 | 0.151 | 16.97 | <.0001 |
| | $X_1^2$ | -0.288 | 0.058 | 11.29 | 0.0009 |
| | $X_2^2$ | -0.163 | 0.044 | 13.91 | 0.0002 |
| | $X_1X_2$ | 0.105 | 0.039 | 7.08 | 0.0083 |
| BM-MSCs | Intercept | -1.874 | 1.142 | 2.69 | 0.1029 |
| | $X_1$ | 3.534 | 1.195 | 8.76 | 0.0035 |
| | $X_2$ | 0.442 | 0.513 | 0.74 | 0.3897 |
| | $X_1^2$ | -0.895 | 0.290 | 9.48 | 0.0024 |
| | $X_2^2$ | -0.064 | 0.148 | 0.18 | 0.6671 |
| | $X_1X_2$ | 0.117 | 0.134 | 0.76 | 0.3855 |

MSCs, mesenchymal stem cells; T-MSCs, tonsil-derived mesenchymal stem cells; BM-MSCs, bone marrow-derived mesenchymal stem cells; $X_1$ , day; $X_2$ , concentration of TGF-β3

In the results, *X*_1_, *X*_2_, 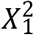, 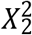, *and X*_1_*X*_2_ were highly significant factors. The analysis of equation (1) depicted that the variables, i.e., days and amount, have positive effects on T-MSCs. Synergistic interactions between days and amount were highly significant (p < 0.05). The results also indicated that the selected quadratic model was adequate in assuming the response variables for the experimental data.

### Three-dimensional response surface plot

MSCs were sensitive to culture time and TGF-β3 concentration, which was consistent with the results presented in Table 5]. Figure 2 depicts the three-dimensional response surface relationship between days and concentration for T-MSCs and BM-MSCs. These data indicate that the appropriate culture days and amount would result in the highest mRNA concentration. Response surface plots as a function of two factors at a time, with maintenance of all other factors at fixed levels, are helpful in understanding both the main and interaction effects of the two factors. In the present study, the interaction effects of the variables and optimal levels of each variable were determined by the response surface graphs. The optimum values drawn from these figures were in close agreement with those obtained by optimizing the regression model equation (1).

There are certain limitations to this study. First, MSCs were stimulated only by a single chemical. Several other proteins can be stimulation candidates for MSCs, although we only referred to previous studies that used TGF-β for tenogenic differentiation of MSCs [8, 22]. Therefore, different approaches would be required for different chemical stimulators. Second, several studies showed that mechanical stretching stimulates MSCs to proliferate and differentiate into tenocytes [23, 24]. Further studies focusing on mechanical stimulation of T-MSCs and BM-MSCs should be performed.

In summary, we demonstrated a protocol for tenogenic differentiation of T-MSCs and BM-MSCs using the DoE approach and showed that this protocol could be less expensive than the standard protocol. In addition, our protocol was optimized with experimental variation. From the DoE approach, tenogenic mRNAs in T-MSCs and BM-MSCs were significantly upregulated at 2.5 days of culture and 2.3 days culture, respectively. In T-MSCs, the peak expression of tenogenic mRNAs was predicted to occur 2.7 ng/mL TGF-β3, whereas no significant TGF-β3 concentration was observed for BM-MSCs. This study suggests the potential of using the DoE approach for optimization of the culture protocol for tenogenesis of MSCs.

